# The IQGAP-related RasGAP IqgC regulates cell-substratum adhesion in *Dictyostelium discoideum*

**DOI:** 10.1101/2024.05.02.592142

**Authors:** Lucija Mijanović, Darija Putar, Lucija Mimica, Sabina Klajn, Vedrana Filić, Igor Weber

**Affiliations:** Ruđer Bošković Institute, Department of Molecular Biology, 10000 Zagreb, Croatia

**Keywords:** cell attachment, DdIQGAP3, amoeboid locomotion, RasG, focal adhesions, cell migration

## Abstract

Proper adhesion of cells to their environment is essential for the normal functioning of single cells and multicellular organisms. To attach to the extracellular matrix (ECM), mammalian cells form integrin adhesion complexes consisting of many proteins that together link the ECM and the actin cytoskeleton. Similar to mammalian cells, the amoeboid cells of the protist *Dictyostelium discoideum* also use multiprotein adhesion complexes to control their attachment to the underlying surface. However, the exact composition of the multiprotein complexes and the signaling pathways involved in the regulation of adhesion in *D. discoideum* have not yet been elucidated. Here we show that the IQGAP-related protein IqgC is important for normal attachment of *D. discoideum* cells to the substratum. Mutant *iqgC*-null cells have impaired adhesion, whereas overexpression of IqgC promotes directional migration. A RasGAP C-terminal (RGCt) domain of IqgC is sufficient for its localization in the ventral adhesion focal complexes, while RasGAP activity of a GAP-related domain (GRD) is additionally required for the proper function of IqgC in adhesion. We identify the small GTPase RapA as a novel direct IqgC interactor and show that IqgC participates in a RapA-regulated signaling pathway targeting the adhesion complexes that include talin A, myosin VII and paxillin B. Based on our results, we propose that IqgC is a positive regulator of adhesion, responsible for the strengthening of ventral adhesion structures and for the temporal control of their subsequent degradation.

## Introduction

Adhesion to the environment is a fundamental property of all living cells. Proper adhesion is critical for numerous biological processes, including motility and phagocytosis of single cells as well as embryonic development, tissue homeostasis and wound healing in multicellular organisms [1]. Adhesion is deregulated in pathological conditions, such as cancer invasion and metastasis [2], cardiomyopathy and atherosclerosis [3]. Adhesion of metazoan cells to their environment is mediated by integrin adhesion complexes (IACs), signaling platforms characterized by the presence of integrins, transmembrane heterodimer proteins that enable coupling of the ECM and the actin cytoskeleton [4]. At integrin sites, the major IAC proteins accumulate in plaques up to 200 nm in size, comprising a layer of signaling proteins such as FAK (focal adhesion kinase) and paxillin, followed by cytoskeletal adaptors such as talin and vinculin, and actin-regulatory proteins such as VASP (vasodilator-stimulated phosphoprotein) and α-actinin [5]. The role of small GTPases such as Rap1 [6], Rac, Cdc42 and Rho [7] is also crucial for the regulation of IAC.

Although the protist *Dictyostelium discoideum* lacks the canonical integrins, other core elements of mammalian IACs are present in this organism, as described in more detail in a recent review [8]. Briefly, several transmembrane proteins have been identified to regulate adhesion, such as Phg1, SadA and SibA [9–12]. Since SibA has partial homology with integrin β-chains, it is thought to be the major adhesion receptor in *D. discoideum* [10], while Phg1 and SadA are thought to play regulatory and accessory roles [13]. Cytoplasmic proteins, such as the FERM (four-point-one, ezrin, radixin, moesin)-domain-containing proteins talin A and B, myosin VII and FrmA-C, have also been associated with the regulation of cell-substratum adhesion in *D. discoideum* [14–19]. Talin A forms a complex with myosin VII [20], which is incorporated into ventral focal adhesion structures followed by the adaptor protein paxillin B [16, 21]. The small GTPase RapA is an essential protein and appears to stimulate adhesion via multiple pathways [22–26]. The small GTPases RasG and RapC have also been shown to positively and negatively influence adhesion, respectively [27–29].

IQGAPs (IQ motif-containing GTPase-activating proteins) are evolutionarily conserved multidomain scaffold proteins that are involved in the regulation of various processes [30]. They consist of six domains: the calponin homology domain (CH), the coiled-coil domain (CC), the WW domain with two conserved tryptophans, the IQ domain (isoleucine–glutamine domain), the GRD domain and the RGCt domain, which together allow IQGAPs to interact with a variety of proteins [31]. The GRDs of IQGAPs show high homology with the RasGAP domains of RasGAP proteins, which deactivate GTPases by simulating GTP hydrolysis, but GRDs are inactived by mutations of crucial residues [31–35]. Instead, GRDs together with RGCt domains mediate the binding of IQGAPs to small Rho GTPases [36–41]. The RGCt domain additionally binds to E-cadherin, β-catenin and PIP2 and contains sites that alter IQGAP function upon phosphorylation [31, 38, 42]. The structure and function of the IQGAP domain are conserved from yeast to humans, and human IQGAP1 has been shown to regulate cell-cell adhesion and migration, among other processes [30, 43]. However, the specific mechanisms of regulation of cell-ECM adhesion by IQGAPs remain largely unexplored. IQGAP1 and IQGAP3 have been identified by proteomics of the adhesome in human K562 and HFF1 cells [44, 45], while IQGAP1 has been shown to be localized in nascent focal complexes and mature focal adhesions [46–48]. It has been suggested that IQGAP1 may be involved in the integration of signaling pathways that regulate adhesion, cytoskeletal remodeling, and phosphoinositide signaling [49, 50].

The genome of *D. discoideum* encodes four IQGAP-related proteins: DGAP1, GAPA, IqgC and IqgD, which are smaller than the mammalian IQGAPs and, with the exception of IqgD, contain only the GRD and RGCt domains in addition to poorly conserved IQ motifs [51]. In contrast to DGAP1 and GAPA, which bind Rho GTPases and have no GAP activity [52–54], IqgC is an atypical IQGAP-related protein. The GRD domain of IqgC retains the conserved residues essential for RasGAP activity, allowing IqgC to function as a RasGAP for the small GTPase RasG [55]. IqgC negatively regulates macroendocytosis [55, 56], contributes to the regulation of cytokinesis [55] and is involved in chemotaxis towards cAMP [57].

To date, IQGAP-related proteins from *D. discoideum* have not been studied in relation to cell adhesion. We found that *iqgC*-null cells adhere poorly to polystyrene cell culture dishes and therefore hypothesized that IqgC positively regulates cell-substratum adhesion. In this work, we use a combination of phenotypic assays, microscopy, genetics and biochemistry to investigate the role of IqgC in this context. We show that IqgC is localized to ventral adhesion foci and that the RGCt domain is necessary and sufficient for its correct localization. Nevertheless, both the RGCt domain and the RasGAP activity of GRD are required to correct the adhesion defect of *iqgC*-null cells. Migration assays show that loss of IqgC differentially affects random motility, chemotaxis to folic acid, and chemotaxis to cAMP. We also show that IqgC interacts with the small GTPase RapA but is not a RapA-directed GAP. We provide evidence that IqgC is involved in the regulation of adhesion via talin A and myosin VII, whereas paxillin B is incorporated into adhesion complexes upstream of IqgC and is required for its proper localization. The small GTPase RasG is not critical for the localization and normal function of IqgC in adhesion, but modulates the rate of IqgC incorporation into adhesion complexes.

Taken together, our results identify IqgC as an important regulator of cell-substratum adhesion and motility of *D. discoideum* cells and indicate that IqgC regulates the maturation and lifespan of adhesion complexes through interaction with RapA and RasG.

## Materials and methods

### Cell culture and transfection

Cells were grown in HL5 culture medium without glucose (Formedium) supplemented with 18 g/l maltose (Sigma) and 50 μg/ml ampicillin and 40 μg/ml streptomycin. Cells were grown at 22°C in polystyrene dishes or shaken in Erlenmeyer flasks at 150 rpm. The parental strain is the axenic *Dictyostelium discoideum* AX2 strain, which is referred to as wild type. Knockout strains in AX2 background used in this work: *iqgC*-null [55], *rasG*-null [58], *talA*-null [14], *paxB*-null [21], *myoVII*-null (this work). Cell lines were transfected by electroporation according to the standard protocol. In brief, 1 × 10^7^ cells were washed with electroporation buffer (10 mM potassium phosphate buffer pH 6.1, 50 mM glucose) and approximately 1 μg of plasmid DNA was electroporated into the cells by pulsing twice at 1000 V with the Xcell gene pulser (Biorad). The cells were incubated with subsequent addition of healing solution (2 mM CaCl_2_, 2 mM MgCl_2_) and transferred to HL5 medium. After 6-18 hours, 10 μg/ml G418 was added and the transformants were maintained in the presence of 10-20 μg/ml G418.

### Vectors

To construct pDM304_YFP-IqgC_mRFP-PaxB/LimEΔCC for the co-expression of fluorescently labeled IqgC and PaxB or LimEΔCC from the same plasmid under the control of identical promoters, pDM304_YFP-IqgC [55] and pDM328_mRFP were used [59]. The coding DNA sequences (CDS) for PaxB and LimEΔCC were amplified by PCR from the cDNA of *D. discoideum* strain AX2. The PaxB fragment was prepared as a BamHI/XbaI insert with cohesive ends compatible with BglII/SpeI sites, and LimEΔCC was prepared as a BglII/SpeI insert. The inserts were ligated into the cut pDM328_mRFP. The entire expression cassette was excised from the pDM328_mRFP-PaxB/LimEΔCC as an NgoMIV fragment and ligated into an NgoMIV-cut pDM304_YFP-IqgC. Previously constructed vectors were used to express the YFP-tagged IqgC variants YFP-IqgC_GRD, YFP-IqgC_RGCt, YFP-IqgC_RGCt-C, YFP-IqgC_ΔGRD, YFP-IqgC_ΔRGCt, YFP-IqgC_Δcentr and YFP-IqgC(R205A) [56].

To perform BiFC experiments, the vectors pDM304_VC-IqgC_VN-RapA(wt/G14V/Q65E/S19N) were prepared from pDM304_VC-IqgC and pDM344_VN [55, 60, 61]. RapA(wt) was amplified by PCR from the previously constructed pGBKT7_RapA(wt), where RapA(wt) was amplified from AX2 cDNA and inserted into pGBKT7 (Matchmaker™ GAL4 Two-Hybrid System 3, Clontech) using BamHI/PstI sites. RapA(G14V), RapA(Q65E) and RapA(S19N) were produced on the RapA(wt) template using the Change-IT^TM^ Multiple Mutation Site Directed Mutagenesis Kit (Affymetrix, USB). All RapA variants were amplified as BamHI/SpeI fragments and fused to the N-terminal part of the Venus fluorescent protein (VN, 1-210 aa) by ligation into the pDM344_VN vector using BglII/SpeI restriction sites. The entire expression cassette was excised with NgoMIV and ligated into the previously constructed plasmid for the expression of IqgC fused to the C-terminal part of Venus (VC, 211-238 aa), pDM304_VC-IqgC [55].

To purify recombinant GST-tagged domains of IqgC for the biochemical experiments, pGEX-6P-1_IqgC [55], pGEX-6P-1_GST-IqgC(GRD) and pGEX-6P-1_GST_IqgC(RGCt) were used [56]. To obtain expression vectors for full-length (FL) and truncated (ΔCAAX) RapA variants with N-terminal HA tags, the sequences for RapA(wt), RapA(Q65E) and RapA(S19N) were amplified as BamHI/SpeI fragments and cloned into the pDM304_HA vector [55]. To purify the RapA protein for the GAP assay, pGEX-6P-1_RapA(wt) was constructed by amplification of RapA(wt) from pDM344_VN-RapA(wt) and insertion into pGEX-6P-1 with BamHI/SalI. pDM304_YFP-RasG(wt) was constructed previously [61]. For pDM304_YFP-RapA(wt), RapA(wt) was amplified from pGBKT7_RapA(wt) as a BamHI/SpeI fragment and inserted into pDM304_YFP using BglII/SpeI sites. pDM304_YFP-RalGDS(RBD) was constructed by amplification of RalGDS(RBD) CDS from HEK293 cDNA and insertion into pDM304_N-YFP using BglII/SpeI sites. All constructed vectors were verified by sequencing. The sequences of all oligonucleotides used in this work are listed in Table S1.

### Generation of *myoVII*-null cells

Cells with deletions in the *myoVII* gene were generated using the CRISPR/Cas9 system according to a previously published protocol [62, 63]. The sgRNAs were designed using Cas Designer [64] and cloned into pTM1544 using Golden Gate Assembly. pTM1544 enables targeted deletions by simultaneous transient expression of Cas9 nickase and two sgRNAs in *D. discoideum* cells. After electroporation of AX2 cells with 10 μg pTM1544_MyoVII, the cells were selected with 10 μg/ml G418. Clonal cell lines were isolated by spreading dilution on *K. aerogenes* lawns on SM agar plates. After colony growth, cells were transferred to 24-well plates filled with G418-free HL5 medium and used for colony PCR to amplify and select the mutated region for successful targeting. The deletion of the target sequence was additionally confirmed by sequencing the target site.

### Detachment experiments

The detachment experiments were performed according to a previously published protocol [11]. Subconfluent cells were seeded one day before the experiment in HL5 medium on 60-mm polystyrene dishes at 1×10^5^ cells/mL. The next day, plates were shaken on an orbital shaker Unimax 1010 (Heidolph) in fresh HL5 medium at 70 rpm for one hour or at 120 rpm for 30 minutes at 22°C. For the first time point (0 minutes), the cells were counted without shaking. After the indicated time, detached cells were collected for counting and attached cells were resuspended and collected. Samples were concentrated by centrifugation and quantified using a Neubauer counting chamber or a CellDrop automated cell counter (DeNovix). The results were expressed as the percentage of cells adhering to the substratum out of the total number of cell.

### Microscopy

For analysis of protein localization in the ventral membrane by TIRF microscopy, subconfluent vegetative cells were seeded on a 35 mm diameter glass bottom dish (MatTek) one day before the experiment. Cells were grown in HL5 medium containing 10 μg/ml G418, which was replaced with LoFlo medium (Formedium) a few hours before imaging. TIRF microscopy was performed using the Dragonfly 505 system (Andor, Oxford Instruments) based on the Nikon Ti2-E inverted microscope, equipped with a CFI Apochromat TIRF 60×C/1.49 oil objective (Nikon) and a Sona 4.2B-6 camera (Andor, Oxford Instruments). For visualization of YFP-tagged fusion proteins by TIRF microscopy, an excitation wavelength of 514 nm and a 538/20 detection filter were used. For YFP and mRFP co-expressor strains, 488- and 561-nm laser lines with 521/38-nm and 594/43 detection filters, respectively, were used. Images of randomly migrating cells were acquired at 1-s intervals with the TIRF penetration depth set to 100 nm. For BiFC, cells were grown in HL5 medium containing 10 – 30 μg/ml G418. Approximately two hours before the experiment, the cells were seeded on a 35 mm diameter glass bottom dish (MatTek) and the HL5 medium was replaced with LoFlo medium. Confocal microscopy was performed using a Leica TCS SP8 X laser scanning microscope (Leica Microsystems) equipped with a HC PL APO CS2 63×/1.40 oil objective. For the BiFC experiments, an excitation of 515 nm and a detection in the range of 525 – 600 nm was used.

### Cell spreading experiments

Vegetative cells were seeded at low density in HL5 medium on glass coverslips and incubated at 22°C in a humid environment. After 45 minutes, images of random positions on the coverslip with cells were captured using a Leica TCS SP8 X confocal microscope (Leica Microsystems) equipped with a HC PL APO CS2 63×/1.40 oil immersion objective in reflection mode to visualize the cell membrane closely apposed to the substratum by RICM. The spreading dynamics of the cells were evaluated by imaging individual cells immediately after plating at an image acquisition rate of 2.58 seconds until spreading was complete and the cells began to migrate.

### Migration and chemotaxis experiments

Random migration experiments were performed as follows: subconfluent vegetative cells were seeded on a 35 mm diameter glass bottom dish (MatTek) at low density in HL5 medium and allowed to rest for one hour prior to the experiment. The day before the experiment, G418 was removed from the medium of IqgC-overexpressing cells. 0.5 mg/ml Texas Red Dextran, 70000 MW (Invitrogen) was added to the HL5 medium immediately before the experiment to facilitate automated tracing. Cells were tracked for one hour with an image acquisition rate of 10 seconds using a Leica TCS SP8 X confocal microscope equipped with a HC PL APO CS2 63×/1.40 objective and a pixel size of 0.48 μm. Excitation wavelengths of 511 and 595 nm with detection ranges of 520-565 and 597-655 nm, respectively, were used to image YFP and Texas Red Dextran. Image processing and migration analysis were performed using the ImageJ TrackMate plug-in [65–67]. Migration tracks were created using DiPer macros for Microsoft Excel [68]. Cell speed was calculated based on the displacement of the cell centroid during the 10-second interval, while directional persistence was calculated as the ratio between the net displacement of the cell and its total trajectory length.

To assess chemotaxis to cAMP, confluent cells were washed twice and starved in Soerenson phosphate buffer for at least 6 hours until streams of cells formed. Aggregation-competent cells were resuspended and seeded at low density on a glass coverslip of the Dunn chamber [69]. After the cells had attached, the chamber was assembled and the outer well was filled with 5 µM cAMP. The chamber was sealed with the hot sealing mixture, which consists of paraffin, beeswax and Vaseline in a 1:1:1 ratio. After a short incubation, the cells were imaged every 10 seconds under a dark-field microscope with a 5× objective [70]. The cell tracks were recorded and analyzed as described for random cell migration. For chemotaxis to folic acid, a chemotaxis assay was performed under agarose according to a previously published protocol [71]. In brief, SM medium containing 1% agarose was poured into 60-mm polystyrene Petri dishes. After polymerization, portions of the gel were cut out to form three parallel wells in the agarose gel (2 mm wide, 39 mm long, and 5 mm apart). 100 µl of 1 mM folic acid was added to the middle well, and approximately 10^6^ cells in 100 µl were seeded into the other two wells. The plate was incubated for 4 hours at 22°C and imaged every 10 seconds with a 5× objective under a dark-field microscope. The detection and analysis of cell traces was performed as described above.

### Biochemical assays

GST-RapA(wt), GST-IqgC_GRD and GST-IqgC_RGCt were produced in the *E. coli* strain Rosetta 2. Expression of the recombinant proteins was induced by 0.75 mM isopropyl-β-D-thiogalactoside (IPTG) overnight at 21°C. Proteins were purified from bacterial lysates by glutathione-agarose affinity chromatography (Thermo Scientific) according to the manufacturer’s instructions. GST-IqgC and GST were purified previously [55]. To test the binding of IqgC and RapA, purified GST-IqgC, GST-IqgC_GRD, and GST-IqgC_RGCt bound to glutathione-agarose beads were used for pull-down assays, while GST served as a negative control. AX2 cells were transfected with constructs expressing HA-RapA (wt/Q65E/S19N) versions of the protein. The expression of the recombinant proteins was checked by Western blotting with monoclonal anti-HA antibodies (Sigma).

Since expression of full-length active proteins could not be detected by Western blotting, we switched to truncated ΔCAAX versions. AX2 cells expressing HA-RapA variants were prepared in lysis buffer (50 mM Tris, pH 7.0, 1 mM EGTA, 40 mM NaCl, 1 mM DTT, 1 mM PMSF, 6.6 mM benzamidine, 0.5 % n-octylpolyoxyethylene, cOmplete™ EDTA free protease inhibitor (Roche)). An immunoblot with a monoclonal anti-HA antibody was performed to check the binding of RapA to GST-tagged proteins. The same *D. discoideum* strains were used for the co-immunoprecipitation test. Protein A-Sepharose CL-4B (GE Healthcare) was incubated with cell lysates and polyclonal anti-IqgC serum [55], and monoclonal anti-HA was used for reciprocal co-immunoprecipitation. The bound proteins were immunoblotted with monoclonal anti-HA and polyclonal anti-IqgC.

The GAP activity of IqgC against RapA was tested using the GTPase-Glo^TM^ assay kit (Promega) according to the manufacturer’s protocol. 1000 ng of previously purified IqgC [55] was added to 1 μM H-Ras (Calbiochem) as a control or to purified GST-RapA(wt) bound to glutathione agarose, and the GTPase reactions were incubated at RT for 2 hours. Luminescence was measured using an Infinite 200 multimode plate reader (Tecan).

### Statistical analysis

The parametric Student’s t-test and the non-parametric Mann-Whitney U-test were used for pairwise comparison of samples. When comparing multiple samples, parametric ANOVA with the Tukey-Kramer post-hoc test and the non-parametric Kruskal-Wallis test followed by the Dunn post-hoc test were used to test for statistical significance. The following significance parameters were used: (*ns*) for p > 0.05, (*) for p < 0.05, (**) for p < 0.001 and (***) for p < 0.0001.

## Results

### IqgC localizes to ventral adhesion foci and is required for proper attachment to the substratum

We found that *iqgC*-null cells are easily washed out of the cell culture dishes during cultivation. To quantitatively test whether IqgC is involved in regulating cell-substratum adhesion, we performed a detachment assay to determine the percentage of cells that remain attached after shaking the dishes on an orbital shaker. The experiment showed that mutant *iqgC*-null cells indeed detached more easily from the polystyrene surface compared to wild-type cells (Fig. 1a). After one hour of shaking, 76.0 ± 1.6% of wild-type cells remained attached, compared to 54.4 ± 4.6% of *iqgC*-null cells (mean ± SEM). This defect was corrected by expressing the recombinant IqgC protein in *iqgC*-null cells (rescue, rsc; 77.9 ± 2.1% of cells remained attached). To determine the localization of IqgC, we expressed YFP-IqgC in wild-type cells and observed the ventral surface of migrating cells with TIRF microscopy. YFP-IqgC was enriched in stationary dot-like structures that remained visible up to the posterior edge of the cell (Fig. 1b). Two classes of discrete punctate structures form on the ventral membrane of *D. discoideum* cells: adhesion foci and F-actin dots [21]. To determine the nature of IqgC-containing structures, we co-expressed YFP-IqgC with mRFP-tagged markers for adhesion complexes, paxillin B (PaxB) [21], or cortical F-actin, LimEΔCC [72]. YFP-IqgC colocalized with mRFP-PaxB (Fig. 1c), but not with mRFP-LimEΔCC (Fig. S1a). The dynamics of recruitment of YFP-IqgC and mRFP-PaxB into the adhesion foci were also similar (Fig. 1d), and the total duration of their fluorescence signals was comparable (YFP-IqgC: 95.0 ± 10.9 s, mRFP-PaxB: 98.0 ± 12.7 s; Fig. 1e), whereas LimEΔCC showed different dynamics (Fig. S1b).

**Fig. 1.**
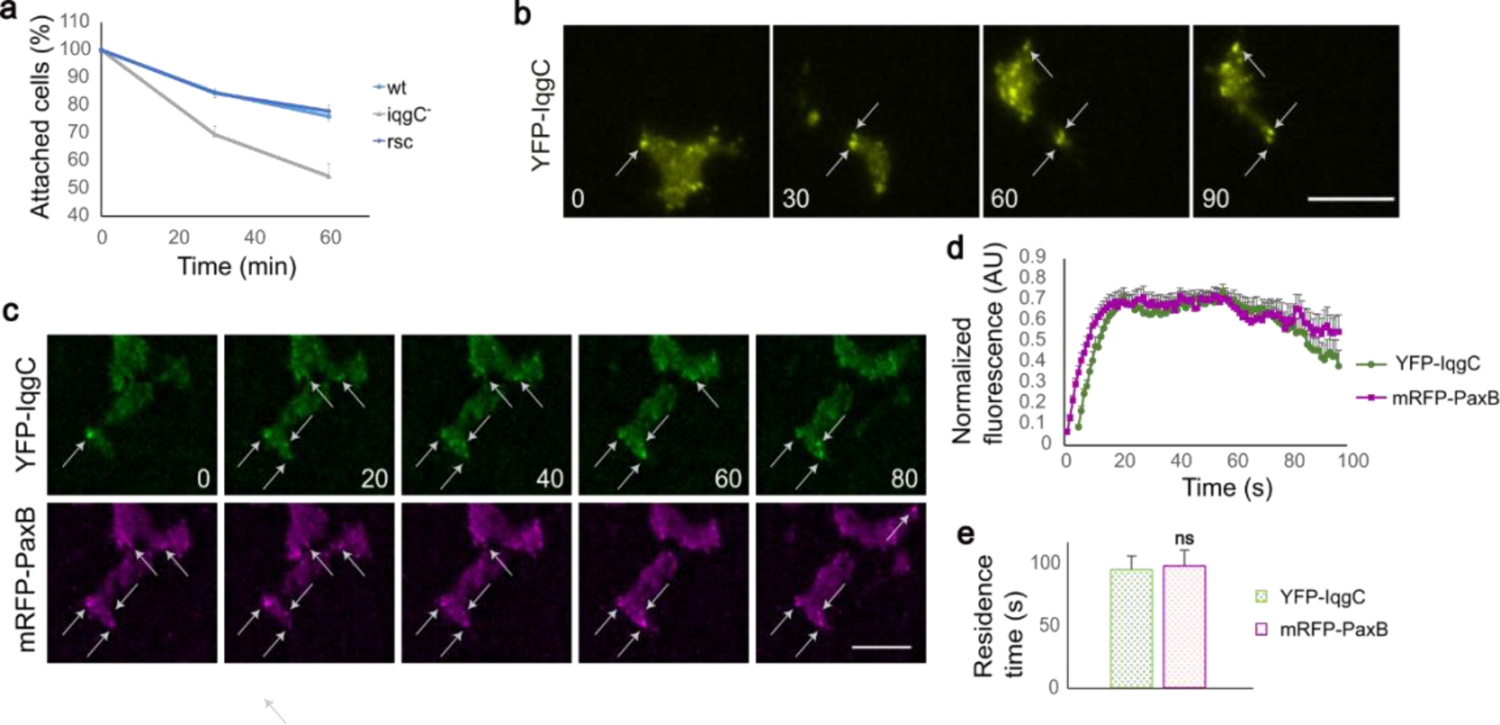
IqgC positively regulates cell-substratum adhesion. **a** Cells lacking IqgC adhere less strongly to the substratum, as shown in the detachment experiment. The number of attached wild-type (wt), *iqgC*-null (iqgC-) and *iqgC*-null cells expressing recombinant IqgC (rescue, rsc) was determined after 60 minutes of shaking (mean ± SEM, n (experiments) ≥ 7). There is a significant difference between wt and iqgC-cells (p < 0.005), but not between wt and rsc (p > 0.7), Mann-Whitney U test. **b** YFP-IqgC localizes to the ventral foci during cell migration, as shown by TIRF microscopy. **c** YFP-IqgC and mRFP-PaxB colocalize at the ventral foci during cell migration. **d** Dynamics of YFP-IqgC and mRFP-PaxB in adhesion foci. **e** The mean residence times of YFP-IqgC and mRFP-PaxB in the adhesion foci are indistinguishable. Data in panels d and e are means ± SEM; n (experiments) = 3; n (foci) = 23; Mann-Whitney U test; ns, not significant. In panels **b** and **c**, the time is given in seconds. Scale bars, 10 µm.

However, mRFP-PaxB was generally detected earlier in adhesion foci than YFP-IqgC (Δt = 4.0 ± 2.2 s). We note that the observed dynamics of paxillin B in adhesion foci and of LimEΔCC in F-actin puncta are consistent with previous reports [21, 73].

### The RGCt domain is necessary and sufficient for the localization of IqgC to ventral adhesion foci

To investigate the importance of the individual IqgC domains for its localization and function in cell-substratum adhesion, we expressed the following fluorescently labeled constructs in wt and *iqgC*-null cells: the single domains, YFP-IqgC_GRD and YFP-IqgC_RGCt; the truncated constructs, YFP-IqgC_ΔGRD, YFP-IqgC_ΔRGCt, and a construct lacking part of the protein between the two domains, YFP-IqgC_Δcentr; a mutant lacking GAP activity, YFP-IqgC(R205A), and the RGCt domain together with the extreme C-terminus, YFP-IqgC_RGCt-C [56]. Cells expressing these proteins were observed by TIRF microscopy as they migrated on a glass surface. Only the constructs encompassing the RGCt domain localized to the adhesion foci (Fig. 2a), but accumulated in the foci at a slightly slower rate than the full-length IqgC (Fig. 2b). Interestingly, the residence time of YFP-IqgC_RGCt in the adhesion foci was shorter (43.0 ± 3.9 s) than that of the full-length protein (87.2 ± 8.6 s), which was partially reversed in the presence of the extreme C-terminus (YFP-IqgC_RGCt-C, 74.0 ± 7.1 s), as shown in Figs. 2b and 2c. Notably, YFP-IqgC_RGCt-C tended to form large aggregates that floated in the cytoplasm (Fig. S2). The construct without GRD domain, YFP-IqgC_ΔGRD, had a longer residence time (134.0 ± 23.0 s), which was apparently not due to the lack of RasGAP activity (YFP-IqgC(R205A), 83.2 ± 9.0 s; Figs. 2b and 2c). Deletion of 88 residues between the GRD and RGCt domains, YFP-IqgC_Δcentr, had no significant effect on residence time in adhesion foci (71.4 ± 12.4 s).

**Fig. 2.**
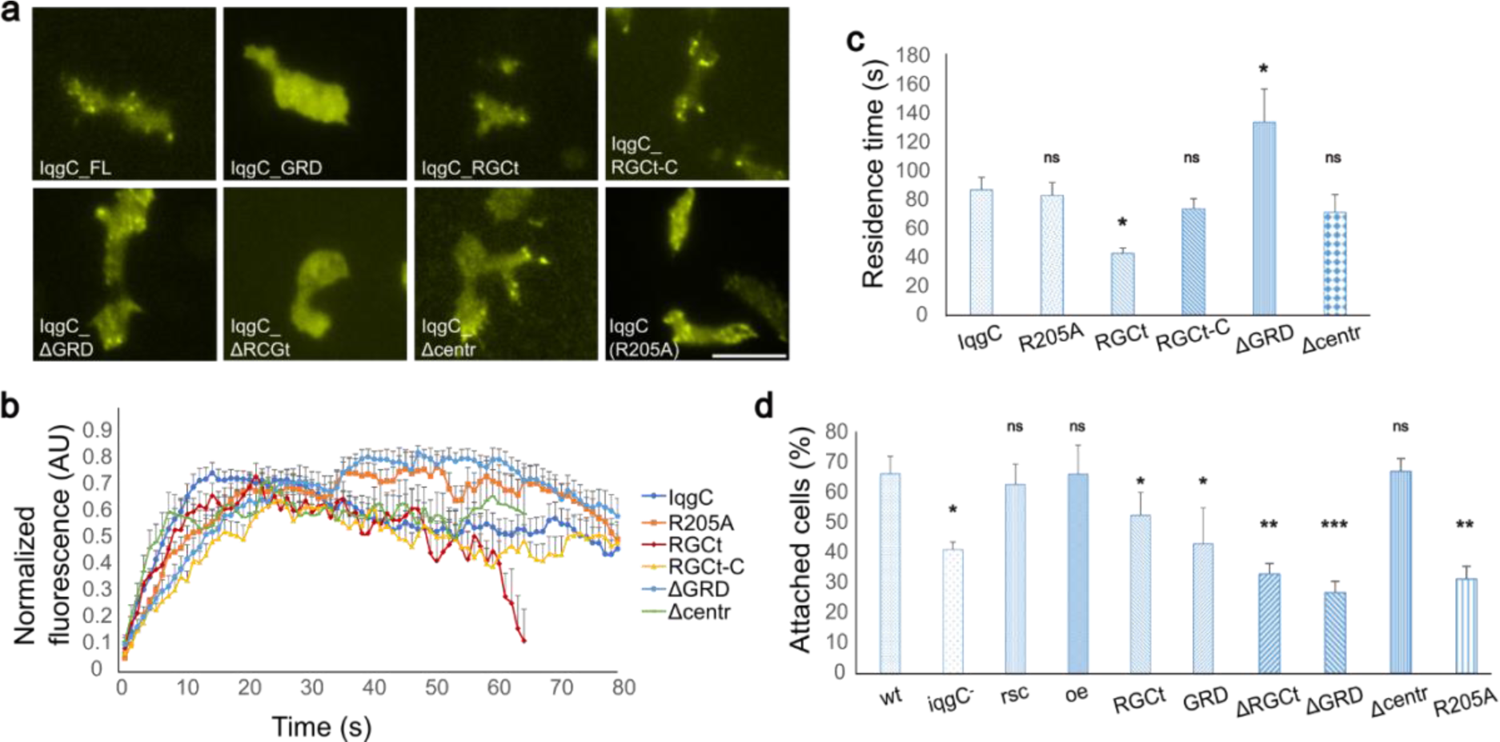
The RGCt domain is necessary and sufficient for the localization of IqgC to adhesion foci, but cannot correct the adhesion defect of *iqgC*-null cells. a TIRF micrographs of wt cells expressing YFP-tagged truncated IqgC variants and the catalytically inactive mutant IqgC(R205A). **b** Time course of incorporation of fluorescently tagged proteins into adhesion foci. **c** Residence times of individual constructs in adhesion foci (mean ± SEM; n (experiments) ≥ 3; n (foci) ≥ 14; ANOVA followed by Tukey-Kramer test). **d** Results of the detachment experiment for wild-type (wt), *iqgC*-null (iqgC-), rescue (rsc) and overexpressor (oe) cells and for *iqgC*-null cells expressing recombinant IqgC variants (mean ± SEM; n (experiments) ≥ 4; ANOVA followed by Tukey-Kramer test). Scale bar, 10 μm.

We next examined detachment of *iqgC*-null cells expressing different IqgC constructs to test for possible rescue of the adhesion phenotype. Under the conditions of our detachment assay, 66.6 ± 5.8% of wild-type cells remained attached, compared to 41.3 ± 2.5% of *iqgC*-null cells and 63.0 ± 6.7% in the rescue strain (Fig. 2d). Additional expression of YFP-IqgC in wt cells (overexpressor, oe) did not alter their detachment (66.5 ± 9.6% of attached cells). Expression of individual RGCt and GRD domains did not rescue the adhesion defect of *iqgC*-null cells, although there was a slight improvement in the case of the RGCt domain (52.7 ± 7.7% for YFP-IqgC_RGCt and 43.2 ± 12.1% for YFP-IqgC_GRD). Interestingly, constructs lacking either the RGCt domain, the GRD domain or RasGAP activity showed a more severe defect than *iqgC*-null cells (32.9 ± 3.5% of cells expressing the YFP-IqgC_ΔRGCt protein remain attached, 26.8 ± 3.8% of cells expressing YFP-IqgC_ΔGRD and 31.2 ± 5.3% of cells expressing YFP-IqgC(R205A)). The only construct that could resolve the adhesion defect of *iqgC*-null cells was YFP-IqgC_Δcentr, as 66.9 ± 4.3% of *iqgC*-null cells expressing this construct remained attached after 60 minutes of shaking.

### *iqgC*-null cells have an enlarged surface in close contact with the substratum

In some *D. discoideum* mutants, a change in adhesion strength is accompanied by a change in the cell area in close contact with the substratum, the contact area, which appears dark when imaged by reflection interference contrast microscopy (RICM). For example, the contact area is smaller in adhesion-disrupted *talA*-null cells [14], whereas the highly adhesive *dymB*-null cells have a larger contact area [74]. We therefore used RICM to test whether the weaker adhesion of *iqgC*-null cells is associated with a smaller contact area. Unexpectedly, *iqgC*-null cells exhibited a larger contact area than wild-type cells (83.3 ± 4.6 μm^2^ vs. 50.0 ± 1.7 μm^2^). This effect was reversed by the overexpression of IqgC (50.5 ± 2.2 μm^2^; Fig. 3a-c). This increase in contact area in *iqgC*-null cells could be partly a consequence of a slight cytokinetic defect in the mutant cells leading to their slightly larger size [55]. However, the dynamics of spreading were similar in wild-type and mutant cells (Fig. 3d).

**Fig. 3.**
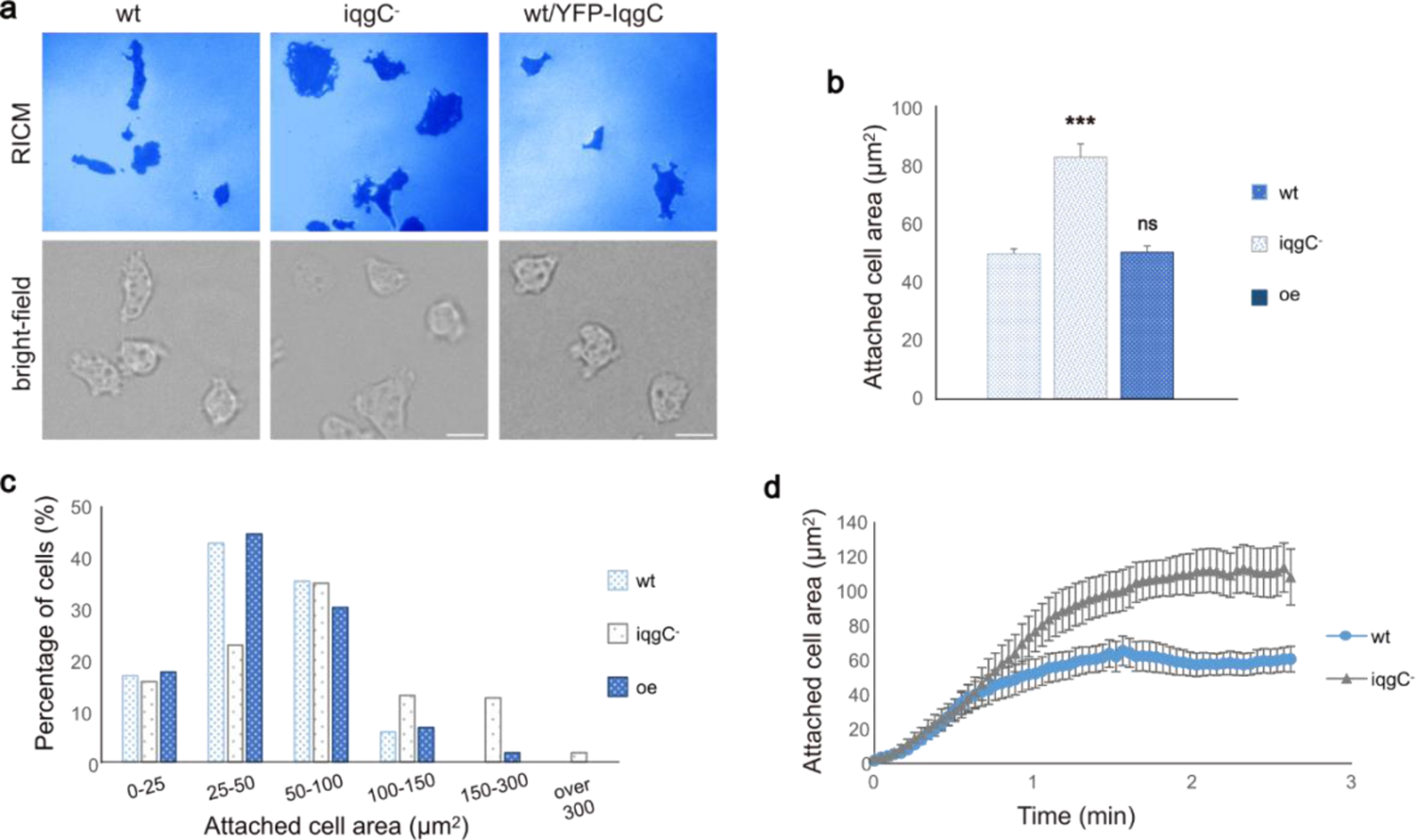
*iqgC*-null cells have a larger contact area with the substratum but their spreading dynamics are similar to wild-type cells. a RICM (upper panel) and bright-field microscopy (lower panel) of wt, *iqgC*-null and oe cells. **b** Dark-appearing membrane in RICM images of wt, *iqgC*- and oe cells (mean ± SEM; n (experiments) = 3; n (cells) ≥ 224; two-tailed t-test). **c** Histogram distribution of the data shown in panel **b**. **d** Spreading kinetics of wt and *iqgC*-cells (mean ± SEM; n (experiments) = 3; n (cells) ≥ 15). Scale bar, 10 μm.

### IqgC is involved in the regulation of random migration and chemotaxis

Since the efficiency of cell migration depends on optimal cell-substratum adhesion, we investigated the speed and directional persistence of cells with different IqgC expression levels. In a random motility assay, both *iqgC*-null and IqgC-overexpressing (oe) cells migrated slightly faster than wild-type cells (Table 1; Fig. 4a-c). In 50 strongly IqgC-overexpressing cells, there was a positive correlation between IqgC expression level and speed (Pearson correlation coefficient equal to 0.689; Fig. 4d). The persistence of *iqgC*-null cells was lower than that of wild-type cells, while the persistence of IqgC-overexpressing cells was higher than that of wild-type cells (Table 1). There was also a positive correlation between IqgC expression level and cell persistence in 50 highly IqgC-overexpressing cells (Pearson correlation coefficient equal to 0.433; Fig. 4e). Interestingly, the highly overexpressing cells were more elongated and maintained the same orientation during migration significantly longer than the cells with low IqgC expression (Fig. S3). Chemotaxis experiments showed that *iqgC*-null cells migrate faster in the cAMP gradient than wild-type cells, exhibiting higher persistence (Table 1). Traces of individual cells from representative experiments are shown in Fig. 4f for wild-type cells and in Fig. 4g for *iqgC*-null cells. Although *iqgC*-null cells migrated more slowly in the folate gradient than wild-type cells, their directional persistencies were comparable (Table 1; Fig. 4h-i).

**Fig. 4.**
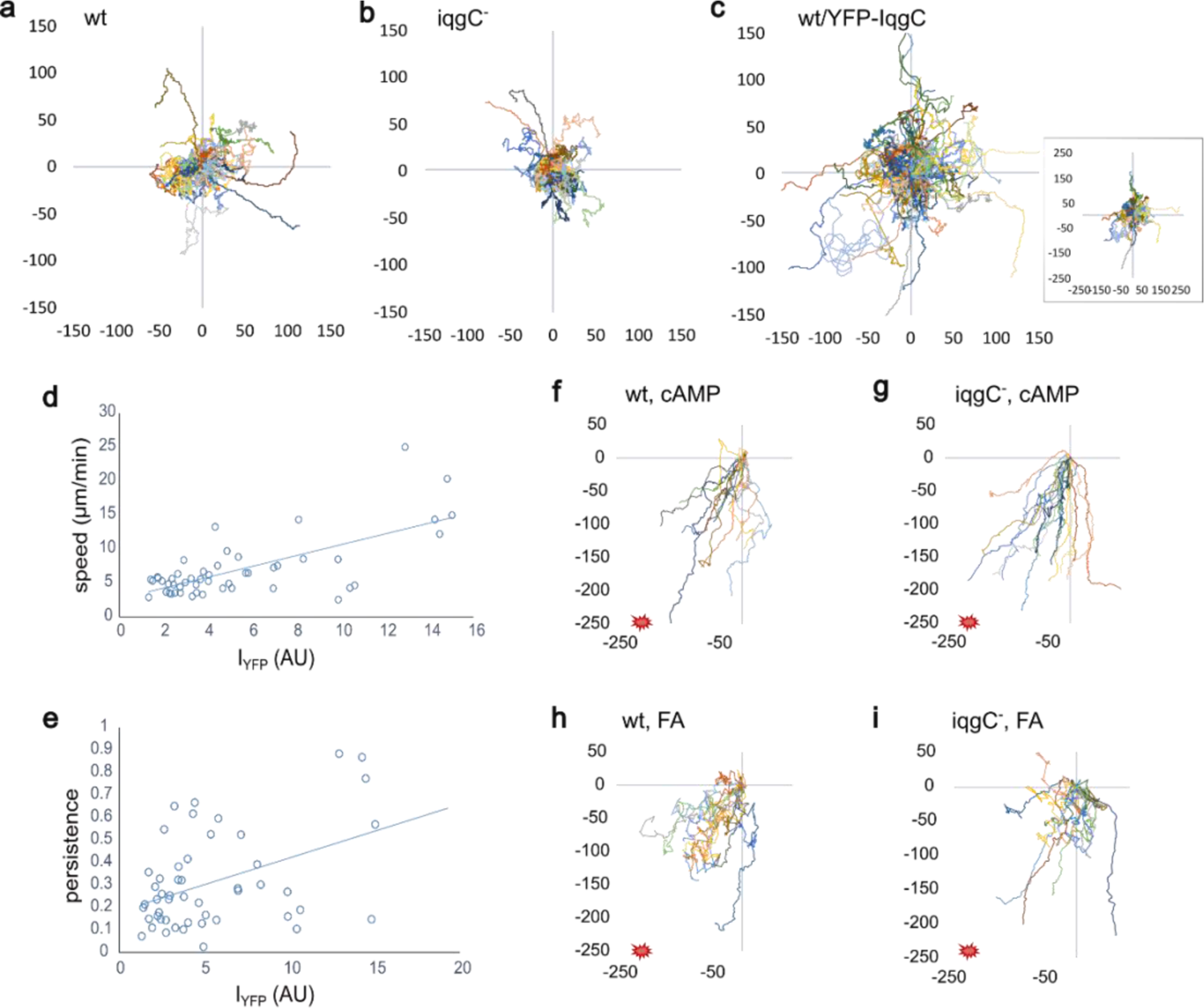
IqgC is involved in the regulation of random migration and chemotaxis. For the random motility assay, vegetative cells were imaged by confocal microscopy in HL5 medium with 0.5 mg/ml Texas Red dextran. **a-c** The trajectories of individual cells during random migration of wild-type (**a**), *iqgC*-null (**b**), and IqgC-overexpressing (oe) (**c**) cells. The inset in panel **c** represents the same experiment as in **c**, but on a larger scale to show the full trajectories of the highly persistent cells. **d-e** Positive correlation of YFP-IqgC expression (I_YFP_) and cell speed (**d**) or persistence (**e**) of 50 highly IqgC-overexpressing cells analyzed. **f-i** Trajectories of single cells during chemotaxis of wild-type and *iqgC*-null cells to cAMP (**f** and **g**) and folate (**h** and **i**) recorded by dark-field microscopy.

**Table 1.** Motility parameters for random migration of vegetative cells in HL5 medium, chemotaxis of aggregation-competent cells to 5 µM cAMP in the Dunn chamber and chemotaxis of vegetative cells to 1 mM folate in the sub-agarose assay. At least 3 experiments per condition were performed. Statistical significance between the mutants and the wild type was tested using the Kruskall-Wallis test (random migration) and the Mann-Whitney U test (chemotaxis).

|  | random migration |  |  | cAMP |  | folate |  |
| --- | --- | --- | --- | --- | --- | --- | --- |
| strain | Wt | *iqgC*<sup>-</sup> | oe | wt | *iqgC*<sup>-</sup> | wt | *iqgC*<sup>-</sup> |
| speed (μm/min) | 5.3 ± 0.2 | 6.0 ± 0.2<sup>ns</sup> | 5.9 ± 0.3<sup>ns</sup> | 8.1 ± 0.2 | 9 ± 0.4* | 12.8 ± 0.02 | 9.4 ± 0.02*** |
| persistence | 0.17 ± 0.02 | 0.11 ± 0.02*** | 0.24 ± 0.02*** | 0.4 ± 0.01 | 0.5 ± 0.02** | 0.34 ± 0.006 | 0.37 ± 0.01<sup>ns</sup> |
| n (cells) | 44 | 44 | 127 | 179 | 60 | 487 | 394 |

### Overexpression of IqgC or RasG abolishes adhesion defects of *rasG*-null or *iqgC*-null cells, respectively

IqgC regulates macroendocytosis by inactivating the small GTPase RasG [55], while both IqgC and RasG act as positive regulators of adhesion [27]. Although these results suggest that IqgC and RasG do not cooperate in the regulation of cell-substratum adhesion, we decided to investigate a possible indirect interplay between the two proteins in this process. First, using the detachment assay, we demonstrated that *rasG*-null cells exhibit an adhesion defect comparable to that of *iqgC-*null cells (27.5 ± 4.5% *iqgC*-null cells and 27.3 ± 2.6% *rasG*-null cells remain attached, compared to 45.3 ± 4.1% wt cells; Fig. 5a). Overexpression of YFP-IqgC in *rasG*-null cells rescued their adhesion defect (43.4 ± 2.3% attached cells), and overexpression of YFP-RasG rescued the adhesion defect in *iqgC*-null cells (41.2 ± 8.2% attached cells). Next, we checked the localization of IqgC in *rasG*-null cells by TIRF microscopy, and detected its accumulation in adhesion foci (Fig. 5b). The residence time of YFP-IqgC in adhesion foci was similar in wild-type and *rasG*-null cells (87.2 ± 8.6 s in wt and 95.1 ± 14.4 s in *rasG*-null cells), but the incorporation of YFP-IqgC was slower in *rasG*-null cells (the fluorescence peak of YFP-IqgC was detected at 27.9 ± 3.8 s in wt versus 47.2 ± 6 s in *rasG*-null cells; Fig. 5c). On the other hand, YFP-RasG was evenly distributed in the ventral membrane of wild-type cells (Fig. S4a). Taken together, these results suggest that RasG and IqgC do not directly influence each other’s role in regulating cell-substratum adhesion, but may belong to converging branches of a signaling pathway that regulates a common effector.

**Fig. 5.**
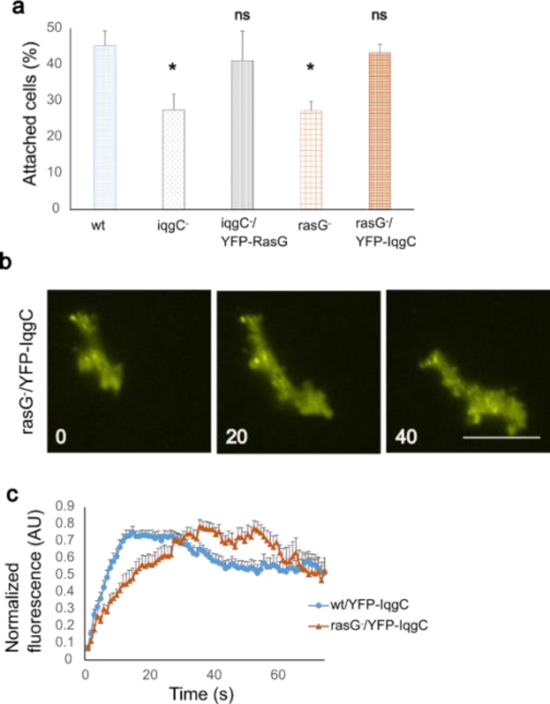
Interdependencies between IqgC and RasG in the regulation of cell-substratum adhesion. **a** Wild-type (wt), *iqgC-*null (iqgC^-^), *rasG-*null (rasG^-^), *iqgC-*null cells expressing RasG (iqgC^-^/YFP-RasG), and *rasG-*null cells expressing IqgC (rasG^-^/YFP-IqgC) were analyzed in a detachment assay (mean ± SEM; n (experiments) ≥ 4; ANOVA followed by Tukey-Kramer test). **b** TIRF microscopy of *rasG-*null cells expressing YFP-IqgC. Time is indicated in seconds. Scale bar, 10 μm. **c** Dynamics of YFP-IqgC incorporation into adhesion foci in wild-type and *rasG-*null cells (mean ± SEM; n (experiments) ≥ 3; n (foci) ≥ 16; Mann-Whitney U test). Scale bar, 10 μm.

### IqgC interacts directly with both active and inactive RapA, but is not a RapA-directed GAP

Next, we decided to search for other potential IqgC interactors based on the published IqgC interactome [55]. The small GTPase RapA caught our attention as it is a known master regulator of cell-substratum adhesion in *D. discoideum* [8, 23]. Therefore, we expressed the wild-type (wt), the constitutively active (Q65E) and the dominant-negative (S19N) HA-tagged RapA variant in wt cells and used the cell lysates to perform pull-down assays with previously purified GST-IqgC [55]. The pull-down assay showed that IqgC interacts with both active and inactive RapA (Fig. 6a). The interaction was further confirmed with the co-immunoprecipitation assay, in which the anti-IqgC antibody was immobilized on protein A-Sepharose beads and used to co-immunoprecipitate RapA with endogenous IqgC from the cell lysates (Fig. 6b). Since we detected weak bands in the negative control, we also performed a reciprocal co-immunoprecipitation assay in which we precipitated the HA-RapA protein with a specific antibody. IqgC was co-precipitated in this assay, confirming an interaction between IqgC and RapA (Fig. 6c). Next, we wanted to investigate which domain of IqgC is responsible for the binding of RapA. IqgC has been shown to bind RasG via its GRD domain [55], but IQGAPs can also bind small GTPases via other domains. For example, IQGAP1 binds Rap1 via its IQ motif [75], while it binds Cdc42 and Rac1 via a multistep mechanism involving RGCt, GRD and the extreme C-terminal part of the protein [38–41]. Pull-down assays with purified GST-tagged RGCt and GRD domains showed that both domains can form a complex with both active and inactive RapA (Fig. 6d-e). HA-RasG was used as a positive control for the IqgC_GRD pull-down [55].

**Fig. 6.**
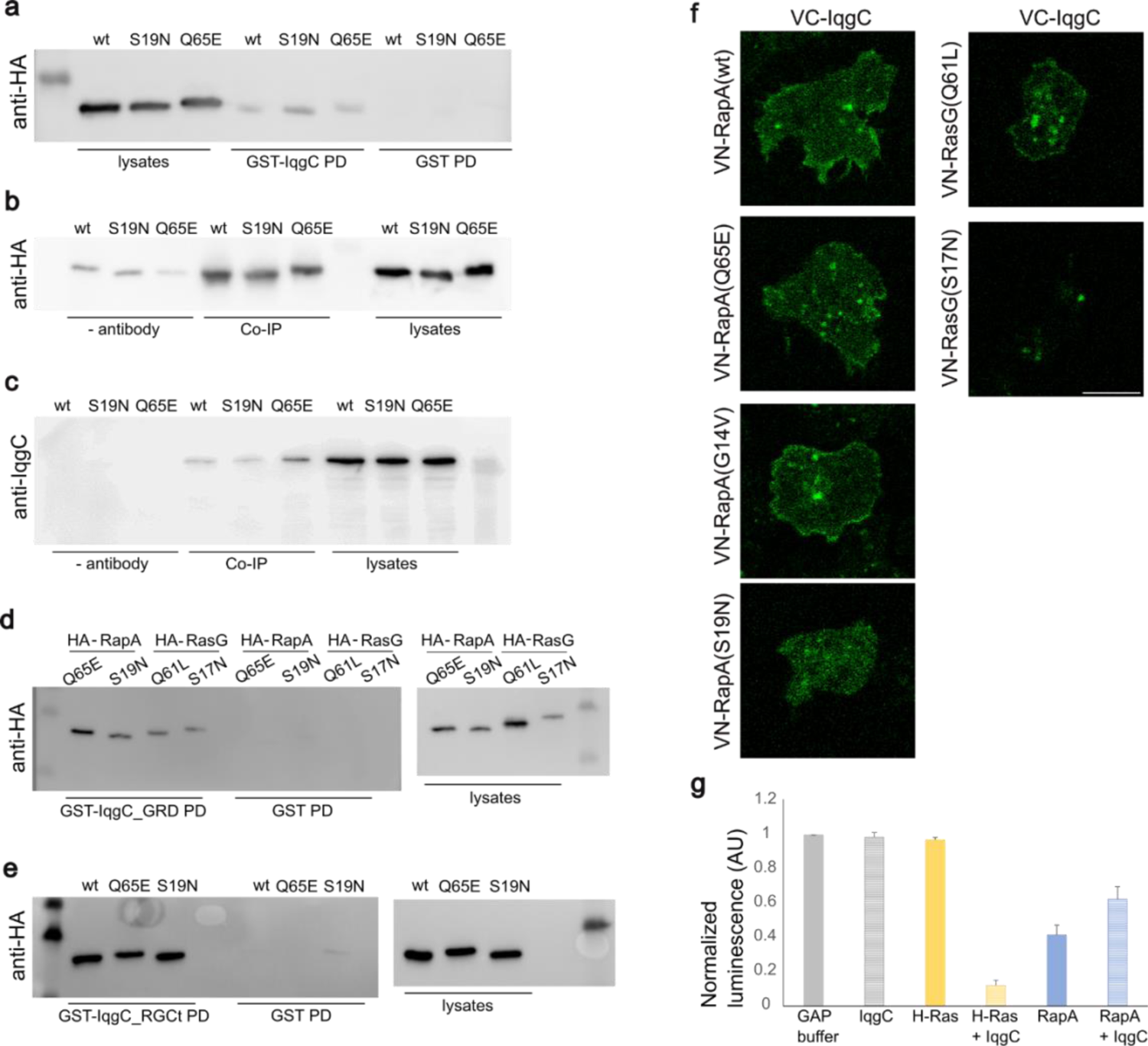
IqgC binds to both active and inactive RapA, but is not a RapA-directed GAP. **a** Pull-down assay (PD) was performed with purified GST-IqgC and lysates from wild-type cells expressing variants of HA-RapA: wild-type (wt), dominant negative (S19N) and constitutively active (Q65E). GST pull-down was used as a negative control. **b** For co-immunoprecipitation (Co-IP), endogenous IqgC was immobilized with an anti-IqgC antibody on protein A-Sepharose and mixed with cell lysates expressing HA-RapA variants. The antibody was omitted for the negative control. **c** Reciprocal Co-IP, in which HA-RapA proteins were immobilized on beads with an anti-HA antibody and used for co-immunoprecipitation of endogenous IqgC. **d-e** Purified IqgC domains, GST-IqgC_GRD in **d** and GST-IqgC_RGCt in **e**, were used for pull-down of HA-RapA variants from cell lysates. Lysates from cells expressing constitutively active (Q61L) and dominant negative (S17N) HA-RasG were used as controls for pull-down with IqgC_GRD. **f** *iqgC*-null cells co-expressing RapA variants fused to the N-terminal part of the Venus fluorescent protein (VN-RapA), and IqgC fused to the C-terminal Venus part (VC-IqgC) were analyzed by confocal microscopy. The interaction of IqgC with all RapA variants *in vivo* is indicated by the fluorescence of the reconstituted Venus protein (left panel). VN-RasG(Q61L) was used as a positive control and VN-RasG(S17N) as a negative control (right panel). Scale bar, 10 μm. **g** A GAP assay was performed to evaluate the potential GAP activity of IqgC against RapA. Purified H-Ras was used as a positive control. When IqgC was added to GST-RapA(wt) immobilized on glutathione agarose (RapA + IqgC), no increase in GTPase activity beyond the intrinsic RapA activity was detected. All values were normalized to the luminescence of the GAP buffer, which was set to 1 (mean ± SEM; n (experiments) = 3). AU = arbitrary units.

To investigate whether IqgC and RapA interact in living *D. discoideum* cells, we decided to use the bimolecular fluorescence complementation (BiFC) assay. We co-expressed the full-length IqgC and RapA variants fused to C- and N-terminal fragments of the fluorescent protein Venus, respectively, in *iqgC*-null cells [60, 61]. While expression of all RapA variants resulted in the appearance of Venus fluorescence, indicating that all RapA variants tested interact with IqgC, the cortical signal was only present in cells expressing the wild-type and constitutively active variants (Fig. 6f). This is consistent with the observation that although RapA is localized both in the cortex and in intracellular structures, RapA activation is restricted to the cell cortex [23].

We used active (Q61L) and inactive (S17N) RasG variants as positive and negative controls, respectively [55]. Although the results of the pull-down, co-IP and BiFC assays did not indicate selective binding of IqgC to activated RapA, we decided to perform a GAP assay to exclude the possibility that IqgC acts as a GAP against RapA. We used a GAP assay in which the luminescence output is inversely correlated with the amount of hydrolyzed GTP. The GTP hydrolysis measured in the GST-RapA-containing sample indicates significant intrinsic GTP-hydrolyzing activity of RapA (Fig. 6g). When IqgC was added to the RapA sample, no additional GTP hydrolysis occurred. On the contrary, there was a slight increase in luminescence (RapA + IqgC), strongly suggesting that IqgC is not a RapA-directed GAP.

### IqgC participates in a RapA-stimulated adhesion pathway

Since we have shown that the small GTPase RapA, a known regulator of adhesion in *D. discoideum*, interacts with IqgC, we decided to investigate the importance of this interaction for cell-substratum adhesion. First, we expressed RapA in *iqgC*-null cells and showed that overexpression of RapA can correct the adhesion defect of the cells (Fig. 7a). In the detachment experiment, 45.3 ± 4.0% of wild-type cells were still attached to the dish after shaking for 60 min, compared with 27.5 ± 4.5% of *iqgC*-null cells and 51.5 ± 5.0% of *iqgC*-null/YFP-RapA cells. A reverse experiment could not be performed because deletion of RapA has been shown to be lethal [23, 25]. Similar to YFP-RasG, YFP-RapA and the fluorescently labeled probe for active RapA, YFP-RalGDS(RBD) [23, 76–78], were evenly distributed in the ventral cell membrane (Fig. S4b), as previously reported [79]. We next investigated the possible interplay between IqgC and other proteins that act downstream of RapA in regulating cell adhesion: talin A, paxillin B and myosin VII [14, 15, 20, 21, 80]. Cells deficient in these proteins (*talA-*, *paxB-* and *myoVII*-null cells) expressing YFP-IqgC were characterized in a detachment experiment. The results show that IqgC was unable to resolve the adhesion defect in any of these strains (Fig. 7b). In this experiment, 79.0 ± 3.4% of wt cells remained attached after 60 minutes of shaking, compared to 53.1 ± 6.6% of *iqgC*-null cells. Significantly fewer *talA*-null and *myoVII*-null cells remained attached (34.6 ± 4.2% and 31.3 ± 3.8%, respectively), and ectopic expression of IqgC failed to correct their adhesion defect (23.1 ± 2.9% for *talA*-null/YFP-IqgC and 34.5 ± 3.6% for *myoVII*-null/YFP-IqgC).

**Fig. 7.**
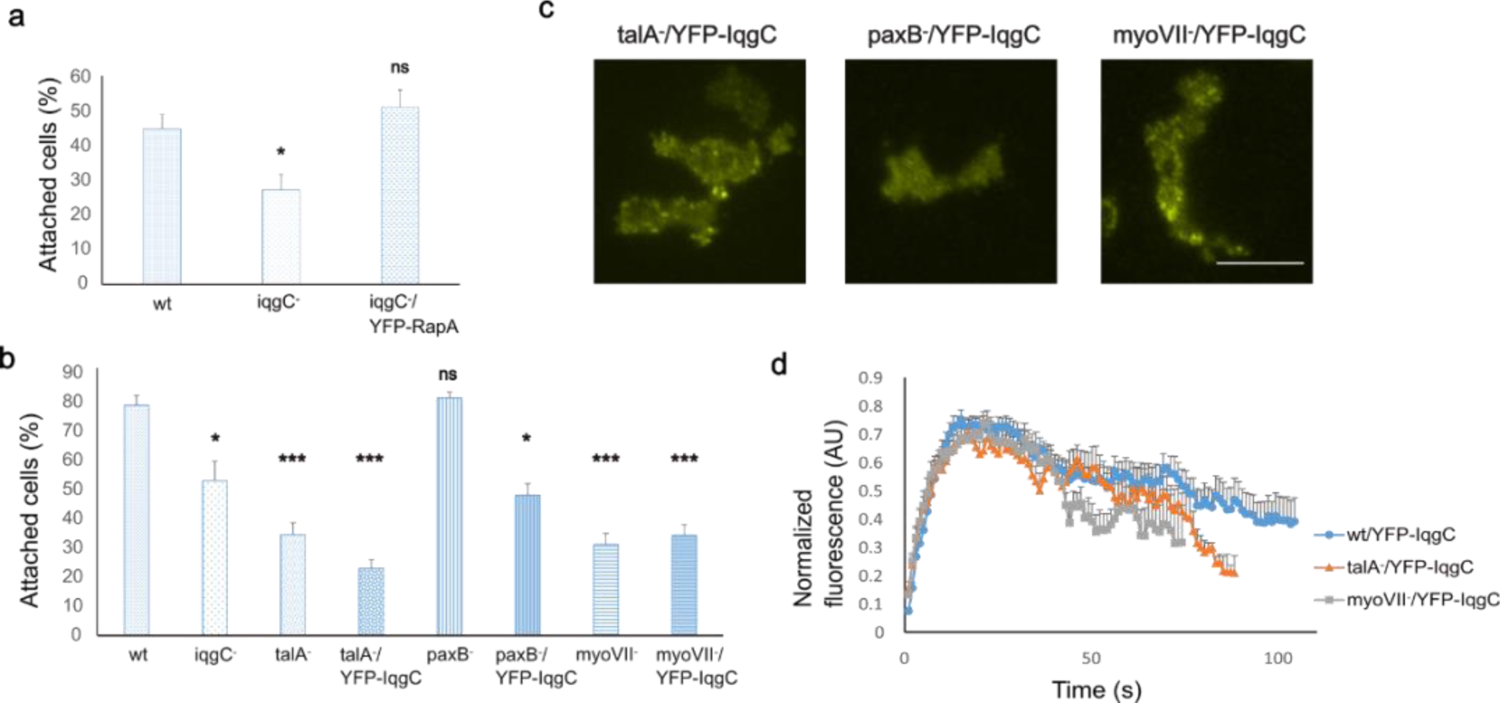
The interplay between IqgC and other proteins involved in RapA-stimulated adhesion. **a** Detachment experiment shows that RapA expression in *iqgC*-null cells (iqgC^-^/YFP-RapA) resolves their adhesion defect (mean ± SEM; ANOVA followed by Tukey Kramer); n(experiments) ≥ 3). **b** IqgC overexpression does not correct the adhesion defect in cells lacking adhesion-regulating proteins downstream of RapA. The detachment experiment was performed with wild-type (wt), *iqgC*-null (iqgC*^-^*), *talA*-null (talA^-^), *paxB*-null (paxB*^-^*) and *myoVII*-null (myoVII*^-^*) cells as well as with the corresponding IqgC-expressing mutants (talA^-^/YFP-IqgC, paxB^-^/YFP-IqgC, myoVII^-^/YFP-IqgC) (mean ± SEM; n (experiments) ≥ 3; ANOVA followed by Tukey Kramer). **c** TIRF microscopy shows that YFP-IqgC is localized in adhesion foci in *talA-*null and *myoVII*-null cells, but not in *paxB*-null cells. **d** Dynamics of YFP-IqgC incorporation into adhesion foci in *talA*-null and *myoVII*-null cells (mean ± SEM; n (experiments) ≥ 3; n (foci) ≥ 28; Kruskal-Wallis test). Scale bar, 10 μm.

In our experiments, *paxB*-null cells adhered normally to the dish surface (81.5 ± 2.0%), but overexpression of IqgC impaired their attachment (48.2 ± 3.9%). Cells from *talA-*, *paxB-* and *myoVII*-null strains expressing YFP-IqgC were analyzed by TIRF microscopy and it was found that YFP-IqgC was not incorporated into adhesion foci only in the absence of PaxB (Fig. 7c). The dynamics of YFP-IqgC incorporation into adhesion foci in *talA*- and *myoVII*-null cells was normal, but the fluorescent signal disappeared from the foci earlier on average than in wild-type cells (Fig. 7d). The main reason for this was the predominance of unstable, short-lived foci, while a small number of foci had a normal duration. In wt/YFP-IqgC cells, YFP-IqgC was present in the adhesion foci for 87.2 ± 8.6 s, in *talA*-null/YFP-IqgC for 49.1 ± 5.2 s, and in the *myoVII*-null/YFP-IqgC strain for 51.5 ± 5.7 s.

Overexpression of PaxB in wild-type cells had no significant effect on their detachment (74.0 ± 5.7% for wt, vs. 84.0 ± 2.0% for wt/YFP-PaxB), but it rescued the adhesion defect of *iqgC*-null cells (50.5 ± 8.3% for *iqgC*-null vs. 70.7 ± 3.0% for *iqgC*-null/YFP-PaxB) (Fig. 8a). YFP-PaxB was localized to adhesion foci in both wild-type and *iqgC*-null cells (Fig. 8b), but we found that YFP-PaxB accumulation and turnover were faster in *iqgC*-null cells than in wt cells (Fig. 8c). YFP-PaxB persisted in adhesion foci of wt cells for an average of 116.9 ± 15.5 s with a maximum intensity at 46.5 ± 4.5 s, whereas in *iqgC*-null cells YFP-PaxB was present in adhesion foci for 50.9 ± 5.9 s with a maximum at 21.1 ± 4.0 s.

**Fig. 8.**
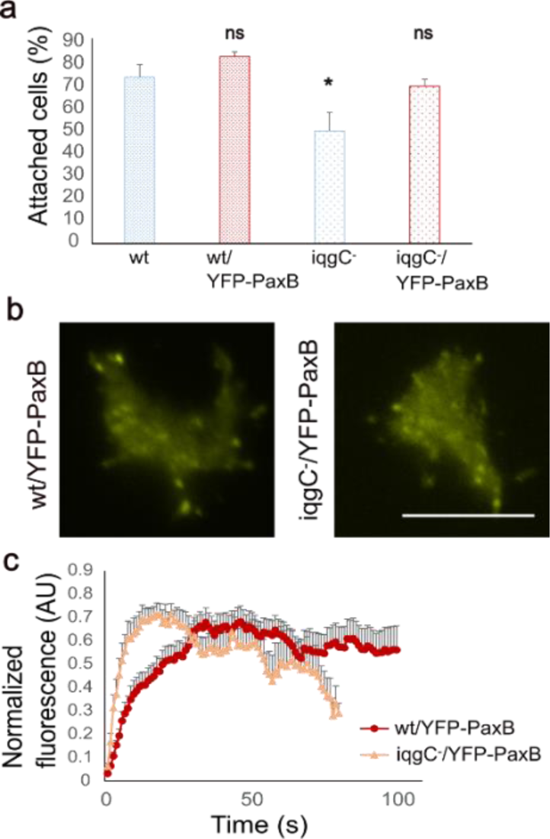
IqgC regulates PaxB dynamics in adhesion foci. **a** Detachment assay shows that overexpression of paxillin B in *iqgC*-null cells (iqgC^-^/YFP-PaxB) resolves their adhesion defect (mean ± SEM; n (experiments) = 3; ANOVA followed by Tukey Kramer). **b** TIRF microscopy shows that YFP-PaxB is localized in adhesion foci in *iqgC*-null cells. **c** Dynamics of YFP-PaxB incorporation into adhesion foci in wt and *iqgC*-null cells (mean ± SEM; n (experiments) = 3; n (foci) ≥ 16; Kruskal-Wallis test). Scale bar, 10 μm.

## Discussion

Although *D. discoideum* amoebae do not possess canonical integrins, they attach to external surfaces by similar mechanisms as cells of metazoans [8]. While the two cell types share the basic components and morphology of the discrete anchoring structures at the ventral cell membrane, the composition and ultrastructure of the adhesion complexes in *D. discoideum* have not yet been elucidated. Here, we identify the IQGAP-related protein IqgC as a novel component of the focal adhesion complexes in *D. discoideum* amoebae. IqgC positively regulates cell-substratum adhesion: *iqgC*-null cells detach more readily from the substratum and YFP-IqgC localizes to paxillin-B-positive adhesion foci. IqgC is an atypical member of the IQGAP family because, unlike other family members in *D. discoideum* and mammals, it has retained RasGAP activity. Recently, it was shown that the GAP activity of IqgC towards RasG is crucial for its role in the downregulation of macroendocytosis [55]. Compared to mammalian IQGAPs, IqgC possesses only GRD and RCGt domains, with the N-terminal motif bearing little resemblance to canonical IQ domains [30]. This domain organization precludes direct binding of IqgC to F-actin via a CH domain, in contrast to mammalian IQGAPs [81, 82]. Interestingly, although human IQGAP1 is localized in nascent adhesions [48] and focal complexes [45–47], its correct localization appears to depend primarily on the IQ motif [48], suggesting that the recruitment of IQGAP proteins to adhesion structures is generally not dependent on their binding to actin.

Our studies on truncated IqgC constructs show that the RGCt domain is sufficient for proper localization, but both RGCt and GRD are essential for the normal function of IqgC in adhesion (Fig. 2). Similarly, interaction of the RGCt domain of IQGAP1 with the adaptor protein β-catenin is sufficient for localization of IQGAP1 to E-cadherin-positive adhesion sites [83, 84], but promotion of strong cell-cell adhesion requires interaction of IQGAP1 with additional proteins [43]. The maturation of cell-cell adhesion complexes involves the interaction of IQGAP1 with the small GTPases Rac1 and Cdc42. Analogously, the formation of functional cell-substratum adhesion complexes in *D. discoideum* could involve the interaction of IqgC with RasG [55] and RapA (Fig. 6). However, these interactions are probably transient, as RapA and RasG are not enriched in the ventral foci (Fig. S4). RasG likely plays a role in the dynamics of IqgC incorporation into adhesion complexes, as recruitment of IqgC to ventral foci is slowed in *rasG*-null cells (Fig. 5c). A possible role of RapA in the formation of focal adhesion complexes is more difficult to assess, as *rapA* knock-out is lethal in *D. discoideum* [23, 25]. However, overexpression of RapA has been shown to stimulate cell-substratum adhesion, in part through Phg2- and talin-mediated mechanisms [22, 24]. Consequently, several RapGAPs of *D. discoideum* have been shown to negatively regulate cell-substratum adhesion: RapGAP1 [79], RapGAPB [85] and RapGAP9 [86], while the RapGEF GbpD positively regulates cell-substratum adhesion [22]. It is therefore not surprising that IqgC, which lacks RapGAP activity and may even inhibit intrinsic RapA GTPase activity *in vitro*, promotes adhesion (Fig. 6g). The possible IqgC-mediated inhibition of RapA GTPase activity is reminiscent of the IQGAP1/2-mediated inhibition of the intrinsic GTPase activities of Cdc42 and Rac1 [36, 37].

Although our results strongly suggest that the presence of paxillin B is crucial for the recruitment of IqgC to the ventral adhesion foci, paxillin B did not appear in the IqgC interactome [55]. However, it is known from other systems that paxillins and IQGAPs can interact. One member of the paxillin family, Hic-5, interacts with mammalian IQGAP1, and is required for the localization of IQGAP1 to invadosomes in HEK 293 cells [87]. In addition, a paxillin-related protein Pxl1p was shown to interact with the IQGAP-related protein Rng2p in the fission yeast *Schizosaccharomyces pombe* [88]. We show that IqgC and paxillin B are sequentially recruited to focal adhesion structures, with IqgC lagging about four seconds behind paxillin B and their net accumulation taking about 20 seconds (Fig. 1d). In agreement with this result, FRAP experiments in MEF cells and zebrafish macrophages showed characteristic turnover times of less than one minute for fluorescently labeled paxillin in focal adhesions and invadosomes [46, 87, 89, 90]. Excessive IQGAP1 accumulation correlated with slower paxillin turnover, elongation of focal adhesions and impaired cell migration [46], suggesting that timely removal of IQGAP1 is important for the dynamics of focal adhesion. Consistent with this, genetic elimination of IqgC resulted in a reduced lifespan of paxillin B-positive adhesion structures (Fig. 7c), indicating a general involvement of IQGAPs in the disassembly of adhesion complexes. Mechanistic details of this regulatory process are not known, but in the case of IqgC, its GRD domain is likely involved, as expression of IqgC_ΔGRD in *iqgC*-null cells prolonged the lifespan of ventral adhesion foci (Fig. 2c).

The fact that overexpression of RasG, RapA, or paxillin B, each of which is a positive regulator of cell-substratum adhesion in *D. discoideum*, resolves the impairment of *iqgC*-null cell adhesion (Figs. 5a, 7a, and 8a) supports the notion that IqgC functions as a scaffolding protein that facilitates interactions within the multiprotein adhesion complex by bringing interacting proteins into close proximity. In the absence of IqgC, the effectiveness of complex formation is reduced, but can be restored by overproduction of other components. Such an effect, in which overexpression of one suppressor gene rescues the mutant phenotype of another gene, is termed a dosage suppression interaction [91]. For example, MLC1 (myosin light chain) acts as a dosage suppressor of a temperature-sensitive mutation in the gene encoding the IQGAP protein of *S. cerevisiae* [92].

Genetic screening in yeast has shown that the vast majority of dosage suppression interactions (∼80%) occur between functionally related genes, but less than half of the gene products involved are linked by a physical protein-protein interaction [93]. It was therefore hypothesized that the products of a gene and its dosage suppressor may be involved in the formation or stabilization of a multiprotein complex without directly interacting with each other. It remains to be clarified whether IqgC and paxillin B interact directly or indirectly within the ventral adhesion complexes in *D. discoideum*.

Since IqgC regulates cell-substratum adhesion, we checked its possible involvement in the regulation of cell spreading and the size of the contact area, the region of the ventral cell membrane that is closely adjacent to the substratum as visualized in RICM. Surprisingly, the absence of IqgC resulted in a larger contact area (Fig. 3).

This phenotype is reminiscent of another strain that exhibits a strong impairment of cell-substratum adhesion, the *sadA*-null mutant strain [11]. Although the contact area of *sadA*-null cells was not measured, these cells exhibited increased size as a consequence of their cytokinesis defect, resulting in a substantial number of multinucleated cells. Recently, it has been shown that *iqgC*-null cells are also multinucleated to a similar extent as *sadA*-null cells and are on average larger than wild-type cells [55]. It is therefore possible that the larger contact area of *iqgC*-null cells is simply a consequence of their larger size.

The slightly increased migration speed of *iqgC*-null cells is probably due to their weaker adhesion to the substratum, which is consistent with the enhanced migration of adhesion-impaired *sadA*-null cells [11] and cells with reduced paxillin B recruitment to focal adhesions [94], as well as with the impaired migration of strongly adherent *frmA*-null cells [16]. This finding also suggests that the constitutive processes involved in cell locomotion, i.e. anterior protrusion and posterior retraction, are not impaired in these cells (Fig. 4). However, deletion of *iqgC* leads to a strong decrease in migration directionality, suggesting a lack of coordination between the constitutive processes that depends on proper cell-substratum adhesion [95]. Consistently, overexpression of IqgC has an opposite effect, increasing directionality. Since detachment under shear stress of IqgC-overexpressing cells was not altered, it is unlikely that adhesion strength *per se* is responsible for the increased persistence of these cells. Instead, improved spatiotemporal coordination of polarized assembly and disassembly of ventral adhesion foci and possibly their increased number is likely responsible. A more detailed analysis of the dynamics of individual foci in *iqgC*-null and overexpressing cells should provide further insights into this question. When *iqgC*-null cells were placed in a cAMP gradient, the orientation defect caused by their deficient adhesion was corrected, likely due to the externally imposed anisotropic stimulus. The persistence of *iqgC*-null cells in chemotaxis was even higher than that of wild-type cells under identical conditions, while the ratio of migration speeds between the two strains remained unchanged, consistent with a previous study [57].

In vegetative *D. discoideum* cells, macropinocytosis competes with migration, as both processes utilize partially identical structural and regulatory proteins [96, 97]. Both macropinosomes and pseudopodia are regulated by PI3K, Ras and Rac activity, and it has been shown that a low local concentration of active Ras leads to the formation of pseudopodia, whereas a high concentration stimulates the induction of macropinocytic cups [97]. We discovered a positive correlation between the level of exogenously expressed IqgC and cell speed (Fig. 4d), although the effect was not significant at the population level, which is due to the high variability of recombinant protein expression. Since RasG is an important promoter of macropinocytosis in *D. discoideum* [98], IqgC-induced deactivation of RasG might shift the balance towards pseudopodia production and favor cell migration over macropinocytosis. In agreement with this interpretation, a negative correlation between the level of IqgC expression and macropinocytosis was recently found [55]. It should be mentioned that both deletion of RasG and exogenous expression of its constitutively active variants lead to slower cell migration [99, 100], which together with our results suggests that tight regulation of Ras signaling is required for adequate control of actin-based cell motility.

The speed of *iqgC*-null cells in chemotaxis to cAMP was increased by about 10% compared to wild-type cells (Table 1), which is similar to the speed difference between vegetative cells of the two strains in random migration and probably has the same adhesion-based origin. However, we note that RasG activity is increased in *iqgC*-null cells, and RasG has previously been shown to be critical for normal chemotaxis to cAMP [101, 102]. Following uniform stimulation of aggregation-competent cells by cAMP, IqgC is transiently recruited to the membrane [55] similar to active Ras [103], while translocation of RapA is slightly delayed [23]. RapA is also activated downstream of RasG in response to cAMP, although the functional role of this signaling pathway is unknown [102]. Since IqgC binds both RasG and RapA, it is reasonable to speculate that IqgC integrates their signaling during chemotaxis. Similarly, the RapGEF GflB of *D. discoideum* has been shown to regulate directional sensing by balancing the activities of Ras, RapA and Rho at the leading edge of chemotaxing cells [104, 105]. In contrast to cAMP-induced chemotaxis, the speed of *iqgC*-null cells was decreased during chemotaxis to folate under agar (Table 1). Although *D. discoideum* amoebae use similar signaling mechanisms in chemotaxis to cAMP and folate [106], they change the mode of locomotion from an actin protrusion-based movement to a predominantly bleb-based migration when covered with a 1% agarose layer [107]. Bleb-based migration is not yet fully understood, but myosin II plays an indispensable role [107–109]. The reduced speed of *iqgC*-null cells in chemotaxis to folate under agar suggests that IqgC may play a role in bleb-based migration. Interestingly, the IqgC interactor RapA promotes the disassembly of myosin II filaments at the leading edge of chemotaxing cells in response to cAMP via the kinase Phg2 [23].

We were able to show that IqgC binds to RapA but has no RapA-directed GAP activity (Fig. 6). Similarly, the human RasGAP p120^GAP^/RASA1 interacts with both H-Ras and Rap1, but only stimulates the GTPase activity of H-Ras [110, 111]. It has therefore been suggested that Rap1 may act as a competitive inhibitor of the GAP-stimulated GTPase activity of Ras [110]. In *D. discoideum*, RasG binds specifically to the GRD of IqgC [56], whereas RapA appears to bind to both the GRD and RGCt domains (Fig. 6). Recently, human IQGAP1 was shown to bind to Cdc42 and Rac1 via three distinct binding sites: GRD, RGCt and the extreme C-terminal part of the protein [38]. According to the proposed mechanism, the RGCt domain establishes an initial high-affinity interaction with the active GTPase, followed by binding of the GRD with low affinity and stabilization of the complex via the extreme C-terminus [38, 39]. Such mechanism has also been proposed for the interaction between Cdc42 and IQGAP2 [40]. A similar multistep mechanism could explain our observation that the RGCt domain alone localizes to adhesion foci but remains there for a much shorter time than the full-length protein or even the RGCt-C construct (Fig. 2c). These results suggest that RGCt is required for recruitment of IqgC to adhesion complexes, possibly mediated by RapA, while the extreme C-terminus appears to stabilize binding. The RasGAP/GRD, on the other hand, could regulate the dissociation of IqgC from adhesion complexes, as the ΔGRD construct remained in the foci for extended periods of time. Whether the binding of RasG to the GRD plays a role in the timely disassembly of adhesion complexes, possibly by displacing RapA, remains to be investigated. Our results also suggest premature disassembly of adhesion complexes in *talA*-null, *myoVII*-null and *iqgC*-null cells, which is associated with reduced adhesion of these cells in the detachment assay (Figs. 7 and 8).

In summary, our results update current concepts on the function and dynamics of ventral focal adhesion sites in *D. discoideum* [8]. The assembly of adhesion complexes is initiated by the binding of talinA, which forms a complex with myosin VII, to the transmembrane adhesion receptor SibA, supported by SadA and Phg2. Paxillin B is then incorporated into the complex, followed by IqgC. These interaction steps are facilitated and regulated by RapA, RasG and other proteins such as FrmA via complex and largely unknown mechanisms that likely resemble the process of IAC assembly in animal cells. Our results suggest that IqgC is an important regulator of the assembly, stability and degradation of focal complexes that control cell-substratum adhesion in *D. discoideum*. The incorporation of IqgC into focal adhesion complexes is mediated by the RGCt domain and regulated by RasG. The GRD domain of IqgC enables its recycling and possibly the timely turnover of adhesion complexes by suppressing Ras signaling. Recently, it has been pointed out that the major limitation to understanding the formation and regulation of focal adhesions during single-cell migration *in vivo* is that there are no model systems in which transient subcellular focal adhesion structures can efficiently form and be easily visualized under high-resolution imaging in the native environment [89, 90]. In this regard, it is worthwhile to investigate the dynamics of focal adhesions in *D. discoideum* amoebae to elucidate the general principles and underlying molecular processes of cell-substratum adhesion [112].

## Statements and Declarations

### Funding

This work has been financed within the Croatian-Swiss Research Program of the Croatian Science Foundation and the Swiss National Science Foundation with funds obtained from the Swiss-Croatian Cooperation Program (IZHRZ0_180584) and supported by funds from the Croatian Academy of Sciences and Arts (10-102/324/66/2021). The work of doctoral student L.M. has been supported by the “Young researchers’ career development project—training of doctoral students” of the Croatian Science Foundation funded by the European Union from the European Social Fund.

### Competing Interests

The authors have no relevant financial or non-financial interests to disclose.

### Author Contributions

Conceptualization: L. Mijanović, V. Filić, I. Weber; Methodology: L. Mijanović, V. Filić, I. Weber; Formal analysis and investigation: L. Mijanović, D. Putar, L. Mimica, S. Klajn; Writing - original draft preparation: L. Mijanović, I. Weber; Writing - review and editing: L. Mijanović, D. Putar, V. Filić, I. Weber; Funding acquisition: L. Mijanović, I. Weber; Resources: V. Filić, I. Weber; Supervision: V. Filić, I. Weber; All authors have read and agreed to the published version of the manuscript.

## Acknowledgments

We thank the Dicty Stock Center (Chicago, IL, USA) for providing *talA*-null and *paxB*-null strains and Tetsuya Muramoto (Toho University: Funabashi, Chiba, JP) for providing the pTM1544 vector.

## Abbreviations

BiFC: bimolecular fluorescence complementation

CC: domain coiled-coil domain

CH: domain calponin homology domain

ECM: extracellular matrix

FAK: focal adhesion kinase

FERM: four-point-one, ezrin, radixin, moesin

GRD: GAP-related domain

IAC: integrin adhesion complex

IQ: domain isoleucine–glutamine domain

IQGAP: IQ motif-containing GTPase-activating protein

NPXY: asparagine – proline – any amino acid – tyrosine

RGCt domain: RasGAP C-terminal domain

RICM: reflection interference contrast microscopy

TIRF: total internal reflection fluorescence

VASP: vasodilator-stimulated phosphoprotein

WW domain: tryptophan-tryptophan domain

## Notes

### Competing Interest Statement

The authors have declared no competing interest.

