## Supplemental Material for "The IQGAP-related RasGAP IqgC regulates cell-substratum adhesion in *Dictyostelium discoideum*"

### Supplementary Information

<u>This file includes:</u>

- Figures S1-S4 with the corresponding captions.
- Table S1 with the corresponding caption.

### Supplementary Figures

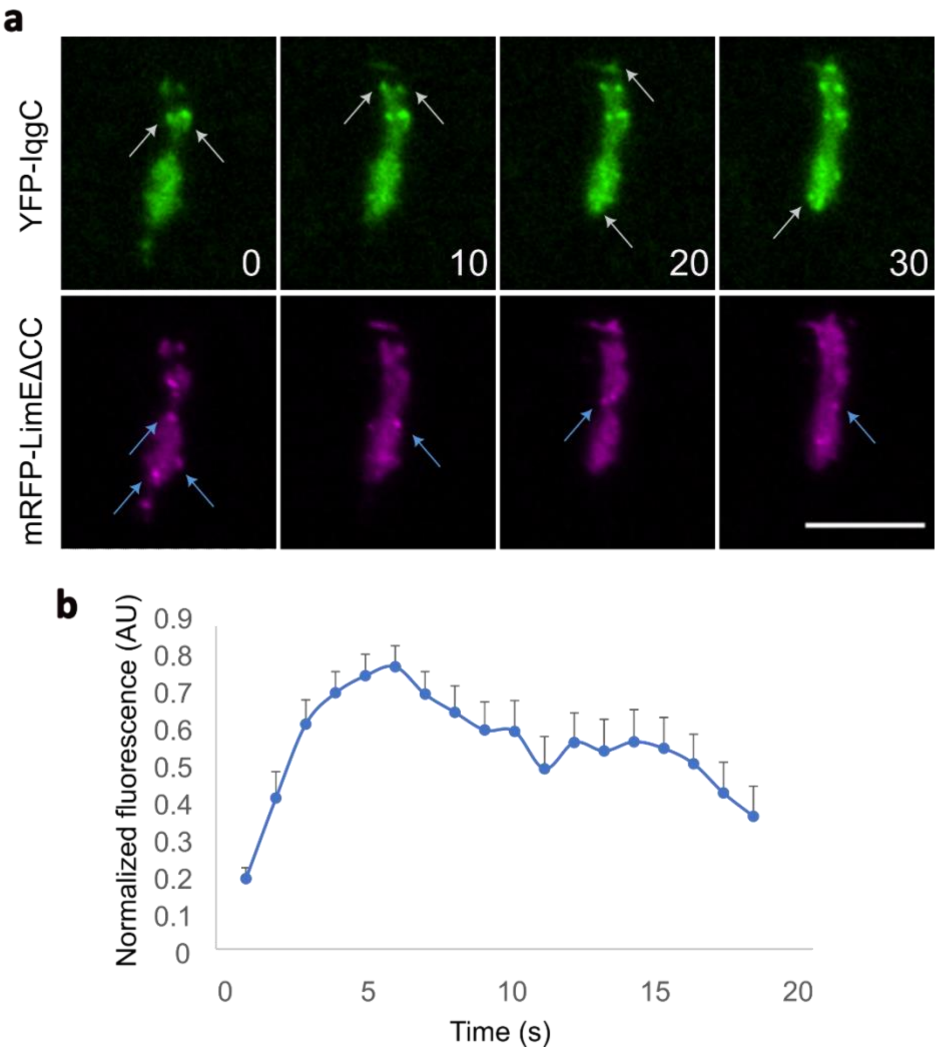


**Fig. S1** IqgC and LimEΔCC do not colocalize in ventral adhesion foci. **a** Dynamics of YFP-IqgC and mRFP-LimEΔCC in ventral foci, as shown by TIRF microscopy. mRFP- LimEΔCC is localized in ventral dot-like structures, which show distinct localization and dynamics in the ventral membrane (cell from Movie 3). **b** The dynamics of mRFP-LimEΔCC in actin dots (n (dots) = 14; n (exp) = 3).

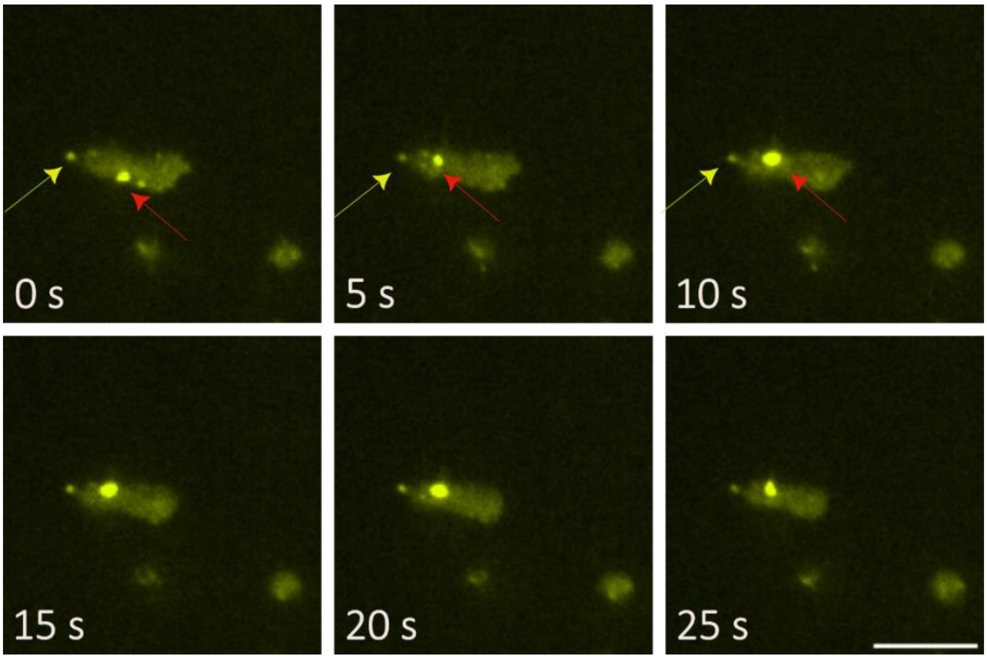


**Fig. S2** A cell expressing YFP-IqgC(RGCt-C) contains a large fluorescent aggregate in the cytoplasm. The aggregates (red arrow) can be distinguished from the adhesion foci (yellow arrow) because the adhesion foci are small and stationary, whereas aggregates are much larger and appear to float around in the cytoplasm. Scale bar, 10 μm.

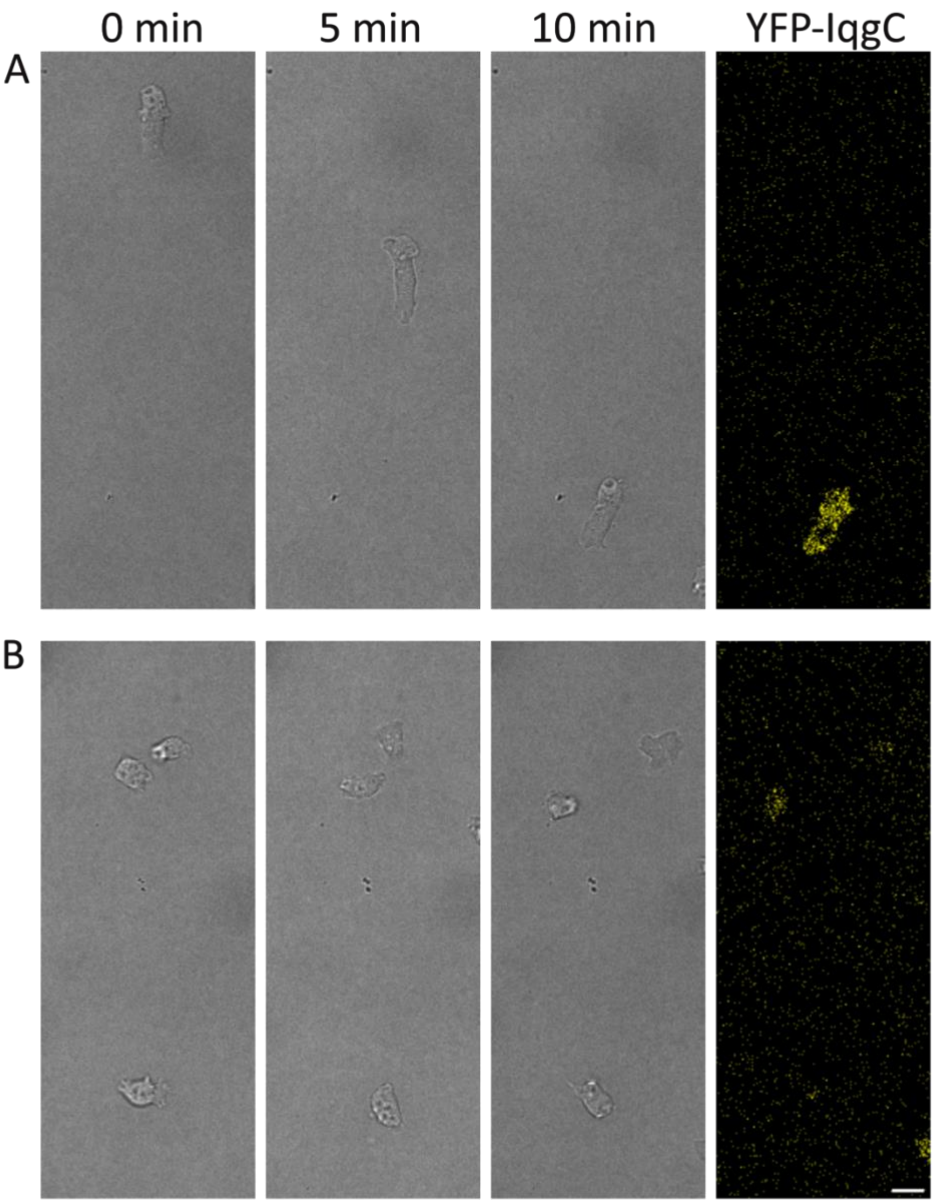


**Fig. S3** Examples of different morphologies and motility of (**a**) cells with higher YFP-IqgC expression that are more elongated and move with higher persistence than (**b**) cells with low YFP-IqgC expression. Cells were observed by confocal microscopy while moving randomly on glass. The brightfield image at 10 minutes is also shown in the YFP fluorescence channel. Scale bar, 10 μm.

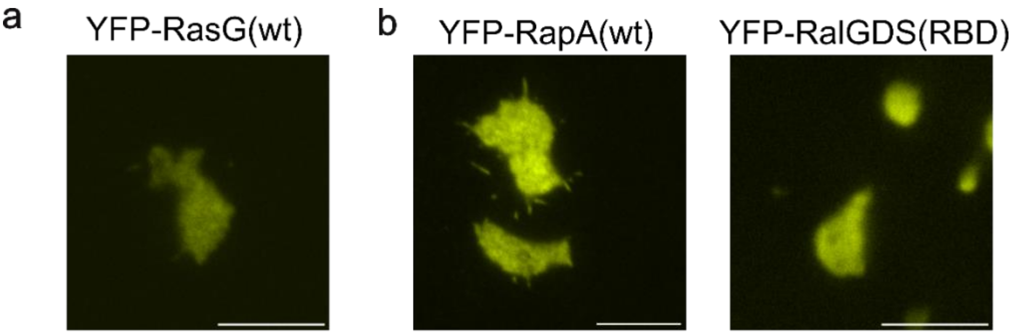


**Fig. S4** Uniform localization of (**a**) YFP-RasG(wt), (**b**) YFP-RapA(wt) and (**c**) a probe for active RapA, YFP-RalGDS(RBD) in the ventral membrane during migration on glass in LoFlo medium, imaged by TIRF microscopy. Scale bar, 10 μm.

### Supplementary Table

**Table S1** List of oligonucleotides used in this study.

| **Usage** | **primer** | **sequence (5' → 3')** |
|---|---|---|
| cloning into pDM304, pDM328 and pDM344 vectors | PaxB_BamHI_F1 | ATTGGATCCATGGCAACAAAAGGATTAAATATG |
| | PaxB_XbaI_R1 | GGCTCTAGATTAAGCAAATAATTTATTATGAC |
| | LimCC-BglII-F1 | ATTAGATCTATGTCTGCTTCAGTTAAATGTG |
| | LimCC-SpeI-R1 | AATACTAGTTTAACCAGTTGGTTGACCATC |
| | RapA_BamHI_F1 | ATTGGATCCATGCCTCTTAGAGAATTCAAAATC |
| | RapA(FL)_SpeI_R1 | AAACTAGTTTACAATAAAGCACATTTTGATTTAGCTTTGCTTG |
| | RapA_SpeI_dCAAX_R1 | ATTACTAGTTTATTTTGATTTAGCTTTGCTTGGTG |
| | RalGDS(RBD)_BglII_F3 | ATTAGATCTATGGACTGCTGTATCATCCGCGTCAG |
| | RalGDS(RBD)_SpeI_R3 | TAAACTAGTTTACCGCTTCTTGAGGACAAAGTC |
| cloning into pGBKT7 vector | RapA_BamHI-F1 | ATTGGATCCAAATGCCTCTTAGAGAATTCAAAATC |
| | RapA_PstI-R1 | TAACTGCAGTTATTTTGATTTAGCTTTGCTTGGTG |
| mutagenesis | RapA_G14V_mut-F1 | CGTCGTTTTAGGTTCAGTTGGTGTAGGTAAATCTGC |
| | RapA_Q65E_mut-F1 | GATACAGCTGGTACTGAAGAATTTACTGCAATGAGAGATC |
| | RapA_S19N_mut_F3 | GGTTCAGGTGGTGTAGGTAAAAATGCTTTGACTGTGC |
| production of *myoVII* null cells | MyoVII_R1_1 | GAGCAAACAATCATTTGTTCGACTTG |
| | MyoVII_R1_2 | TAAACAAGTCGAACAAATGATTGTTT |
| | MyoVII_F1_1 | AGCATCTTTAGCAAAAGCATTATA |
| | MyoVII_F1_2 | AAACTATAATGCTTTTGCTAAAGA |
| | MyoVII_seq_fw | CTCATGTTACCTCATATTTTCG |
| | MyoVII_seq_rev | TCCAATACACCAATAAATGTTGAAT |
| cloning into pGEX-6P-1 vectors | RapA(FL)_SalI_R1 | TAAGTCGACTTACAATAAAGCACATTTTGATTTAGCTTTGC |
| | GST_N-IqgC_Bam-F1 | ATTGGATCCGATCATTATGGACAATTATTTTATTTAC |
| | GST_N-IqgC_Sal-R1 | AATGTCGACTTAGAAACGATTTGTGAGTTTC |
| | GST_C-IqgC_Bam-F1 | ATTGGATCCGTAGATAACATTAAAAAGATATTG |
| | GST_C-IqgC_Sal-R1 | AATGTCGACTTATCTATCTTCTGGTAC |
